# Pathogenic *MAST3* variants in the STK domain are associated with epilepsy

**DOI:** 10.1101/2021.03.09.434675

**Authors:** Egidio Spinelli, Kyle R Christensen, Emily Bryant, Amy Schneider, Jennifer Rakotomamonjy, Alison M Muir, Jessica Giannelli, Rebecca O Littlejohn, Elizabeth R Roeder, Berkley Schmidt, William G Wilson, Elysa J Marco, Kazuhiro Iwama, Satoko Kumada, Tiziano Pisano, Carmen Barba, Eva H Brilstra, Richard H van Jaarsveld, Naomichi Matsumoto, Lance H Rodan, Kirsty McWalter, Renzo Guerrini, Ingrid E Scheffer, Heather C Mefford, Simone Mandelstam, Linda Laux, John J Millichap, Alicia Guemez-Gamboa, Angus C Nairn, Gemma L Carvill

**Affiliations:** Epilepsy Center and Division of Neurology, Ann & Robert H. Lurie Children’s Hospital of Chicago, Chicago, Illinois, USA; Department of Psychiatry, Yale School of Medicine, Connecticut Mental Health Center, New Haven, Connecticut, USA; Division of Genetics, Birth Defects and Metabolism, Ann and Robert H. Lurie Children’s Hospital of Chicago, Chicago, Illinois, USA; Epilepsy Research Centre, Department of Medicine, Austin Health, The University of Melbourne, Heidelberg, Victoria, Australia; Department of Physiology, Northwestern University Feinberg School of Medicine, Chicago, Illinois, USA; Division of Genetic Medicine, Department of Pediatrics, University of Washington, Seattle, Washington, USA; Department of Molecular and Human Genetics, Baylor College of Medicine, Houston, Texas, USA; Department of Pediatrics, Baylor College of Medicine, San Antonio, Texas, USA; Division of Medical Genetics, University of Virginia, Charlottesville, Virginia, USA; Department of Neurology, University of California, San Francisco, California, USA; Department of Pediatric Neurology, Cortica Healthcare, San Rafael, California, USA; Department of Human Genetics, Yokohama City University Graduate School of Medicine, Yokohama, Japan; Department of Neuropediatrics, Tokyo Metropolitan Neurological Hospital, Tokyo, Japan; Neuroscience Department, Children’s Hospital A. Meyer-University of Florence; Genetics Department, University Medical Centre Utrecht, The Netherlands; Department of Neurology and Division of Genetics and Genomics, Boston Children’s Hospital; GeneDx, Gaithersburg, Maryland, USA; Department of Pediatrics and Radiology, University of Melbourne, Melbourne, Victoria, Australia; Department of Medical Imaging, Royal Children’s Hospital of Melbourne, Melbourne, Victoria, Australia; Ken and Ruth Davee Department of Neurology, Northwestern University Feinberg School of Medicine, Chicago, Illinois, USA; Department of Pediatrics, Northwestern University Feinberg School of Medicine, Chicago, Illinois, USA; Department of Pharmacology, Northwestern University Feinberg School of Medicine, Chicago, Illinois, USA

## Abstract

**Objective:** The MAST family of microtubule-associated serine-threonine kinases (STK) have distinct expression patterns in the developing and mature human and mouse brain. To date, only *MAST1* has been associated with neurological disease, with *de novo* variants in individuals with a neurodevelopmental disorder, including a mega corpus callosum.

**Methods:** Using exome sequencing we identify *MAST3* missense variants in individuals with epilepsy. We also assess the effect of these variants on the ability of MAST3 to phosphorylate the target gene product ARPP-16 in HEK293T cells.

**Results:** We identify *de novo* missense variants in the STK domain in 11 individuals, including two recurrent variants p.G510S (n=5) and p.G515S (n=3). All 11 individuals had Developmental and epileptic encephalopathy, with 8 having normal development prior to seizure onset at < 2 years of age. All patients developed multiple seizures types, while 9/11 had seizures triggered by fever and 9/11 had drug-resistant seizures. *In vitro* analysis of HEK293T cells transfected with *MAST3* cDNA carrying a subset of these patient-specific missense variants demonstrated variable but generally lower expression, with concomitant increased phosphorylation of the MAST3 target, ARPP-16, compared to wildtype. These findings suggest the patient-specific variants may confer MAST3 gain-of-function. Moreover, single-nuclei RNA sequencing and immunohistochemistry shows that *MAST3* expression is restricted to excitatory neurons in the cortex late in prenatal development and postnatally.

**Interpretation:** In summary, we describe *MAST3* as a novel epilepsy-associated gene with a potential gain-of-function pathogenic mechanism that may be primarily restricted to excitatory neurons in the cortex.

## Introduction

Developmental and epileptic encephalopathies (DEEs) encompass a group of disorders featuring frequent epileptic activity and developmental impairment. Development plateaus or regresses when seizures are worse but can also occur independent of seizures, and are thus attributable to the underlying cause alone ^1, 2^. DEE causes can be varied, but a genetic etiology is known or suspected in most cases without metabolic or structural causes. The highest yield for genetic testing is in the early onset DEEs where up to 80% of patients have a known underlying genetic etiology ^3^. Both phenotypic and genetic heterogeneity are standard features of DEEs; age of onset of seizures and developmental delay varies and patients have a spectrum of intellectual disability severity and comorbidities. Furthermore, an individual with one type of DEE can evolve to another, and variants in the same gene can lead to a spectrum of DEEs ^1^. In patients with DEEs, the goal of genetic investigations is not only to end the diagnostic odyssey for families but ideally, to identify a potentially actionable target that could lead to precision medicine therapies to help control seizures, rescue the underlying developmental impairments, and ideally arrest further progression. To this end, high-throughput sequencing approaches to gene discovery are an integral part of precision medicine in epilepsy ^4^.

Despite the high diagnostic rate of patients with DEEs, many patients still lack a genetic diagnosis, likely at least in part to pathogenic variants in undiscovered genes. Here we describe variants in *MAST3*, a <u>m</u>icrotubule-<u>a</u>ssociated <u>s</u>erine <u>t</u>hreonine kinase <u>3</u>. This gene is a member of a family of serine threonine kinases (MAST1-4 and MAST-like) that is predominantly expressed in the human cortex, hippocampus, and striatum ^5^. Although MAST3 may play a role in inflammatory bowel disease ^6^, it has not been implicated in a neurological disorder, and its role in the brain is mostly unknown. Pathogenic variants in the *MAST3* homolog, *MAST1*, have been identified in individuals with developmental brain abnormalities that include a particularly striking mega-corpus callosum along with cerebellar hypoplasia and cortical malformations. Individuals also present with a spectrum of developmental impairments including poor speech and a subset with epilepsy ^7^.

As part of an international collaboration, we identified 11 individuals from 4 continents with DEEs and *de novo* variants in the serine/threonine kinase (STK) domain of MAST3. *In vitro* modeling of a subset of these variants showed a potential gain of function effect on MAST3 kinase activity. We propose *MAST3* as a novel cause of DEEs and implicate a second member of the MAST family in neurodevelopmental disorders though with distinct phenotypes, likely related to gene expression timing and/or localization during neurodevelopment.

## Materials and Methods

### Study Participants

All patients were identified by either clinical or research investigations to determine the underlying etiology of their DEE. Patients were ascertained through personal communication or GeneMatcher ^8^. Exome sequencing was performed by a commercial clinical laboratory in the USA or Europe (Patients 1, 2, 6, 7, 10, 11) ^9^, in research laboratories/in-house sequencing facilities (Patients 3, 4) or by exome sequencing of the proband by the Epi25 Collaborative (Epi25 Collaborative, in press) (Patients 5, 9). The variant of patient 8 was identified by targeted resequencing in a research laboratory using single-molecule molecular inversion probes designed to cover the exons +5bp into the intron ^10^. Variant allele frequencies were assessed using GnomAD and TopMed (see list of URLs). All variants were classified according to the American College of Medical Genetics and Genomics (ACMG) guidelines for variant interpretation ^11^. This study was approved by the local institutional review board at each site and all patients and parental or legal guardians provided consent for this study and publication of clinical and genetic data. Missense Tolerance Ratios ^12^ were plotted with ggplot2, combined annotation dependent depletion (CADD), and polyphen scores were generated by variant effect predictor. Clinical histories were analyzed and seizure types and epilepsy syndromes were classified according to the International League Against Epilepsy classification criteria ^13^

### MAST3 expression in the developing human and mouse brain

To determine the expression of MAST3 in the mature and developing human brain we analyzed RNA-seq and single nucleus-RNA-seq data from the Allen Cell Types Database and the Allen Developing Human Brain Database. In addition to RNA expression analysis, we performed immunohistochemistry in 11-week-old human cerebral organoids generated with the STEMdiff cerebral organoid kit (STEMCELL Technologies), and C57BL/6J mouse brains (Jackson laboratory, Stock No:000664) at E14.5 and E16.5. Cerebral organoids and mouse brains were fixed in 4% paraformaldehyde, then cryoprotected in 30% sucrose until sunk. Samples were frozen in an ethanol/dry-ice bath and 10 to 14-micron thick sections were cut with a cryostat (Leica). Sections were permeabilized and blocked for 1-hour at room temperature with 0.3% Triton X-100 and 10% normal donkey serum, respectively, and then overnight at 4°C with the following primary antibodies: MAST3 (Novus Biologicals, Cat # NBP1-82993), CUX1 (Santa Cruz Biotechnology, sc-514008), SATB2 (Abcam, ab51502), TBR1 (Proteintech Group, 66564-1-Ig) and CTIP2 (Abcam, ab18465). After a 2-hour incubation at room temperature with secondary antibodies (Donkey anti-rabbit AlexaFluor 594, anti-rat AlexaFluor 488, anti-mouse AlexaFluor 647), slides were counterstained with Hoechst 33342 (ThermoFisher Scientific) and mounted with ProLong Gold Antifade (ThermoFisher Scientific). Images were acquired with a Nikon A1R laser scanning confocal microscope.

### Analysis of effect of MAST3 variants on phosphorylation of ARPP-16 in HEK293 cells

To assess their kinase activity, MAST3 wild-type and a subset of variants were individually expressed in human embryonic kidney (HEK) 293T cells (ATCC CRL-3216) with the established MAST3 substrate protein ARPP-16 ^14, 15^. All MAST3 constructs were cloned into the pCMV plasmid backbone and fused to a 6x histidine tag on the C-terminus. MAST3 mutations identified in this study, as well as a kinase-dead control (K396H) ^16^,were generated by single-site mutagenesis (Agilent 210518) according to the manufacturer’s instructions. ARPP-16 was expressed under the control of a CMV promoter and fused to a C-terminus hemagglutinin (HA) tag. HEK 293T cells (300,000/well) were plated in 12-well plates and grown in DMEM (Thermo 11995-065) supplemented with 5% FBS and 1% penicillin/streptomycin in a humidified incubator at 37°C and 5% CO_2_ one day prior to transfection. Cells were co-transfected with 250-750 ng each of pCMV MAST3 and pCMV ARPP-16 DNA using Lipofectamine 2000 (Invitrogen 52887) according to the manufacturer’s instructions and incubated overnight. A pCMV GFP construct was used in place of MAST3 for the control well expressing only ARPP-16. The next day, cells were collected in lysis buffer (TBS + 1% Triton X-100) containing protease inhibitors (Complete Mini EDTA-Free, Roche 04693159001), briefly sonicated, and centrifuged at 14,000 x g for 5 min at 4°C. Supernatants were transferred to a new tube and protein concentration measured using the bicinchoninic acid (BCA) assay (Pierce 23225). Protein concentrations were adjusted to 1 ug/uL, and 15 ug was used for analysis by western blot. Samples were mixed with Laemmli sample buffer, denatured at 98°C for 5 min, then separated by SDS-PAGE using Criterion 4-20% Tris-HCl gels (BioRad 3450033). Proteins were transferred to nitrocellulose membranes and probed with HA-tag antibody (1:5000, mouse monoclonal, CST 2367S), MAST3 antibody (1:2000, rabbit polyclonal, Bioworld BS5790), or pS46 ARPP-16 antibody (RU1102, rabbit polyclonal, 1:3000) ^14, 15^. Antibodies were diluted in 2% non-fat dry milk in TBS and incubated overnight at 4°C. The next day, blots were incubated for 1 hour at room temperature with IRDye 680RD goat-anti-rabbit (1:20,000, LiCor Biosciences 926-68071) and IRDye 800CW goat-anti-mouse (1:20,000, LiCor Biosciences 926-32210). Blots were imaged using a LiCor Odyssey system. Band signal intensities were analyzed using ImageJ software and the pS46 ARPP-16 band intensities were normalized to the MAST3 band intensities.

## Results

### Genetic characterization of individuals with MAST3 variants

Twelve individuals with variants in *MAST3* were identified; 3 females and 9 males with a median age of 10.5 years (range 7 to 44 years). 11 individuals carried variants in the Serine Threonine kinase (STK) domain (Table 1, Fig 1A) and one outside this domain. In this study we focus on the variants in the STK domain, all of whom exhibit DEE as described below. The twelfth individual is a 3 year-old boy with a diagnosis of Autism Spectrum Disorder (ASD), but with no history of seizures. He has a a similarly affected brother. We identified a *de novo* c.1963T>C, p.F655L variant (CADD = 24.3 and polyphen = 0.604) in the proband only.

**Table 1.**
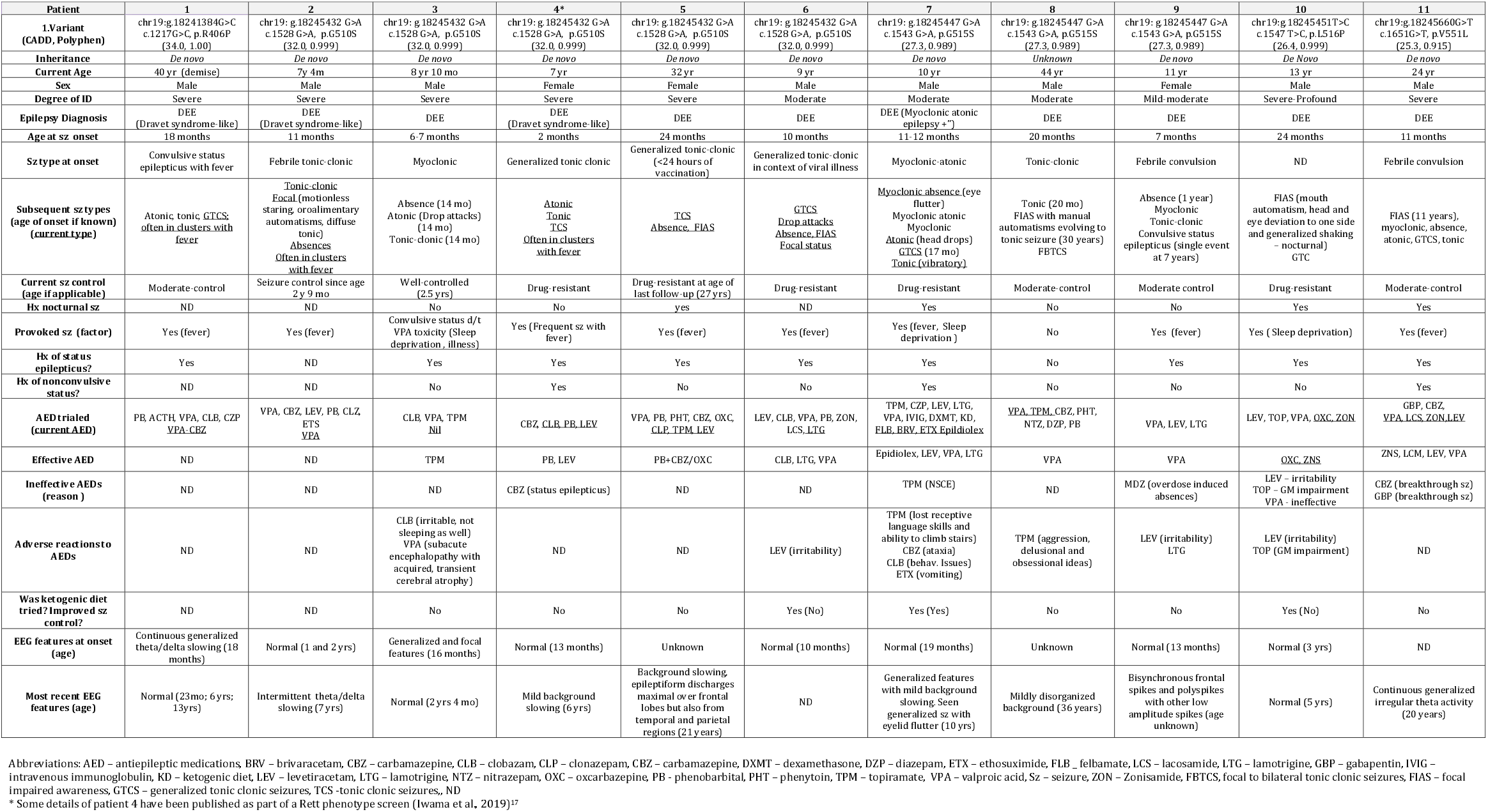
Genetic details and seizure-specific clinical features for individuals with *MAST3* STK domain *de novo* variants

**Figure 1.**
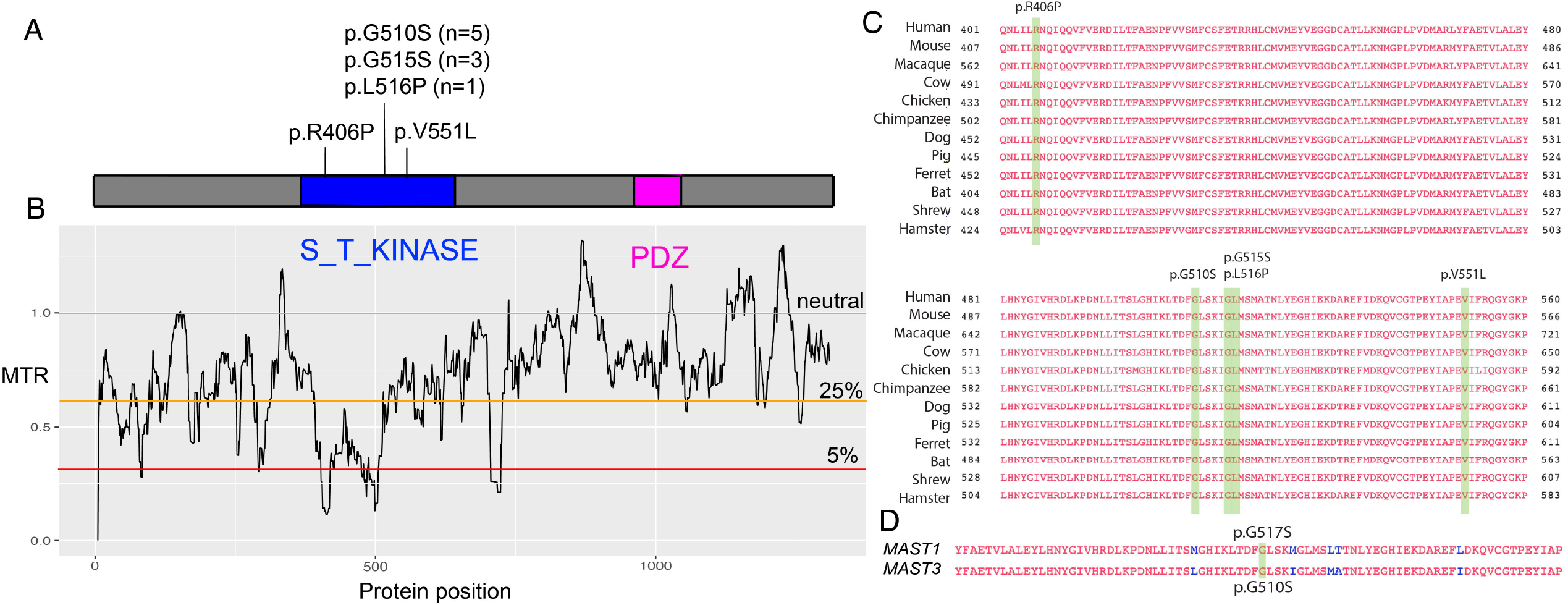
*MAST3* patient-specific variants occur at conserved sites that are intolerant to genetic variation. **(A)**. *MAST3* patient-specific variants (number of individuals in parenthesis) in relation to ST kinase (amino acids 367-640) and PDZ domains (amino acids 958-1038). (**B**) Corresponding graph with missense tolerance ratio (MTR, y-axis) and protein position (x-axis). The tolerance ratio measures cDNA intolerance to missense variants with 5% representing the cutoff for more intolerant segments ^12^. Regions of the ST kinase domain which are intolerant to variations portions of the ST kinase domain overlap with the distribution of pathogenic variants. (**C**) Multispecies alignment of the STK domain harboring the MAST3 patient variants (highlighted in green) show all pathogenic variants are highly conserved across 11 species (identical/similar amino acids in red) (**D**) Alignment of the MAST1 and MAST3 protein sequences. The serine threonine kinase (STK) domain has high homology and the MAST1 p. G517S and MAST3 p.G510S (highlighted in green) occur at the same position in the STK domain. Overall, these protein sequences have 61% identity.

Language development was normal until a regression at 15-18 months leading to both receptive and expressive language delays. Similarly, there was a regression in social development leading to poor eye contact, smile and joint attention. Given that this variant was associated with an atypical presentation as compared to other individuals in this cohort, and the variant was outside of the STK domain, we interpret this variant as a variant of uncertain significance (VUS).

Six unique missense variants were identified in 11 individuals occurring in the STK domain and two variants were recurrent (Fig 1A): c.1528G>A, p.G510S (patients 2-6) and c.1543G>A, p.G515S (patients 7-9). Three additional individuals had unique missense variants in the STK domain (patient 1: c.1217G>C, p.R406P; patient 10: c.1547T>C, p.L516P; patient 11: c.1651G>T, p.V551L).

Of the variants in the STK domain, 10/11 arose *de novo*, while segregation analysis is pending in proband 8 with the recurrent p.G515S variant. All variants were highly conserved (Fig 1C and D) and were predicted to be deleterious or damaging by *in silico* tools (CADD and polyphen respectively) (Table 1) and were absent in controls (GnomAD or TOPMed). *MAST3* is intolerant to variation with a loss-of-function observed/expected upper bound fraction (LOEUF) of 0.6 for missense variation (range 0-1 denoting intolerant to tolerant to missense variation). Moreover, the STK domain is highly conserved and intolerant to variation, with missense tolerance ratio (MTR) scores all below the 25^th^ percentile (Fig 1B) ^12^.

### Epilepsy phenotype of patients with MAST3 STK domain variants

All 11 patients had DEE, including three with a Dravet-like phenotype (Table 1). All patients had seizure onset at or before 2 years of age (median: 11 months, range: 2 to 24 months), with 7/11 patients having seizure onset before 12 months of age. The first seizure was a febrile convulsion, in 4 patients, an afebrile, tonic-clonic seizure in 4, myoclonic seizure in 1 and a myoclonic-atonic seizure in 1 patient (Table 1).

All patients developed multiple seizure types, including atonic (3), myoclonic (2) myoclonic absence with eye flutter, myoclonic-atonic (1), tonic (5), generalized tonic-clonic with or without fever (11) and focal with impaired awareness (6). Seizures evolved to include other types in most patients, and these included atonic, myoclonic, myoclonic absence with eye flutter, myoclonic, atonic, tonic, tonic-clonic, and focal with impaired awareness. All patients experienced generalized tonic-clonic seizures with and without fevers (n = 11). Ten of 11 patients reported provoking factors for seizures (such as illness or fever and sleep deprivation). Eight patients experienced episodes of convulsive status epilepticus and 3 of non-convulsive status epilepticus.

Two patients were seizure free at their most recent follow up: patient 3, now 8 years old, has been seizure free since age 2.5 years and is on no medications while patient 2, now seven years old, has been seizure-free since 2 years nine months on valproate alone. The remaining 9 patients had drug-resistant seizures and were taking between 1-3 (mean: 2) antiepileptic medication with a variety of pharmacological targets represented (Table 1); 3 patients are currently taking sodium channel blockers. Antiepileptic medications that elicited some improvement in seizure control included valproic acid (5), topiramate (1), zonisamide (2), phenobarbital (1), felbamate, oxcarbazepine (1), lamotrigine (1), lacosamide (1), levetiracetam (2), briavaracetam, clobazam and cannabidiol. The ketogenic diet was trialed in 3 patients with improvement in seizure control for patient 7 but not for patient 6 or 10.

Initial EEGs performed within 12 months of seizure onset were available for 8/11 patients; 6 were reported as normal whilst background slowing was reported in patient 1 and generalized and focal abnormalities were reported in patient 3. Follow up EEGs were available for 9/11 patients; 3 were reported as normal whilst 4 patients had background disorganized or slowing, while patient 9 had bisynchronous frontal spikes and polyspikes and patient 7 had generalized seizures with eyelid flutters captured on EEG.

### Neurodevelopmental phenotype of patients with MAST3 STK pathogenic variants

Early development was normal in all 8 individuals with data available (Table 2). All individuals except patient 5, experienced developmental regression or plateau before 2 years of age (mean: 16 months, range: 11-24 months) coinciding with seizure in 6 patients and status epilepticus in 5. Data was unavailable for patient 5. Developmental regression most commonly affected language (7) followed by gross motor (4), social (1), fine motor (1) and toilet training (1). The majority of patients (10/11) are described as non-verbal or with limited speech with expressive speech affected more than receptive speech. Eight patients exhibited delayed gross motor milestones with walking attained at a mean age of 17.5 months (range 14-31 months). Intellectual disability was severe in 6 individuals, moderate in 3, mild-moderate in patient 9 and severe-profound in patient 10 (data not available for patient 5).

**Table 2.**
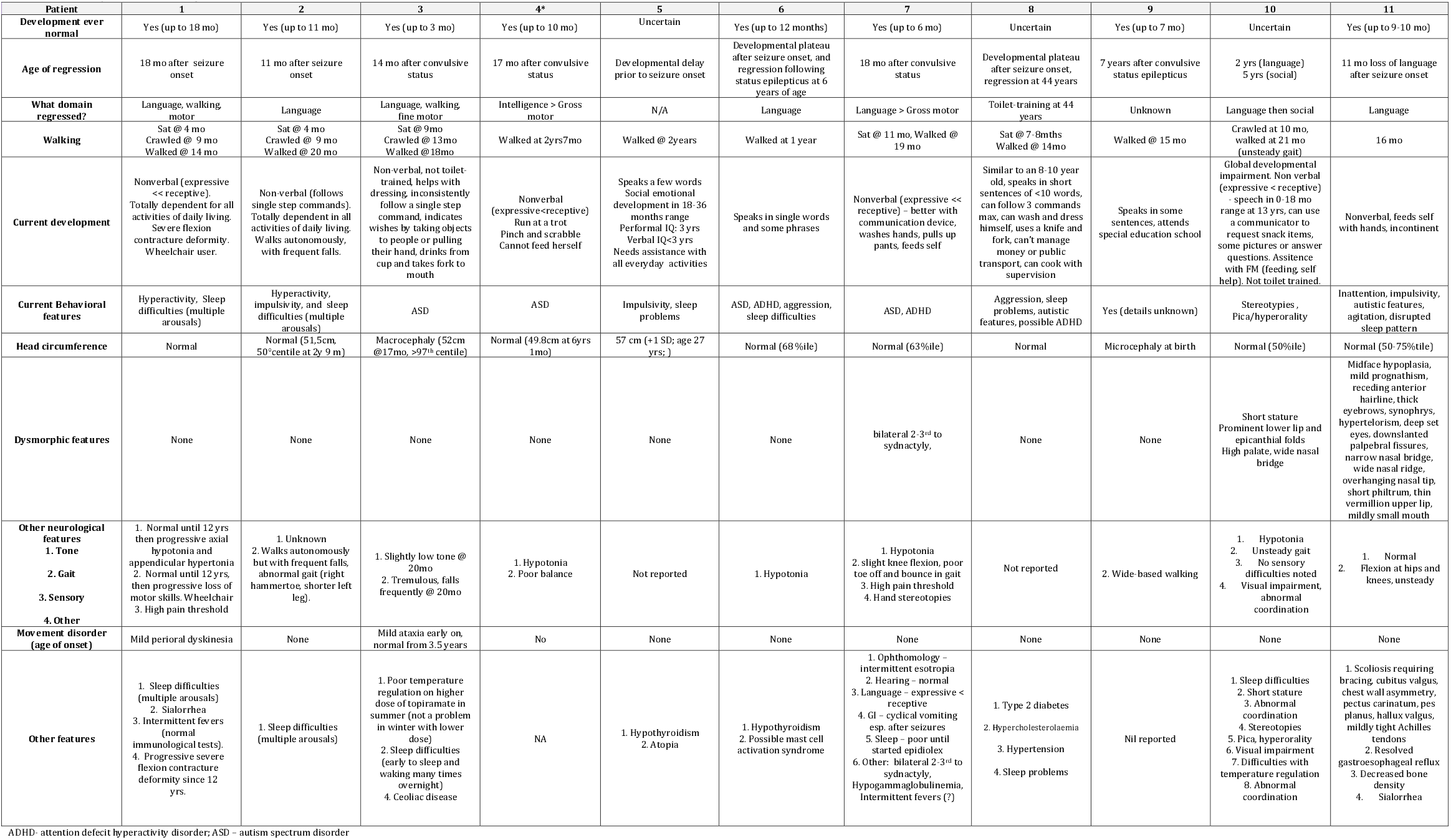
Developmental History and Other Features for individuals with *MAST3* STK domain *de novo* variants

Nine patients were normocephalic whilst patient 3 had macrocephaly and patient 9 had microcephaly. Three of 11 patients had dysmorphic features; patients 10 and 11 had dysmorphic facies, patient 10 also had short stature and patient 7 had 2-3^rd^ toe syndactyly. Six patients had hypotonia with patient 1 developing progressive axial hypotonia and appendicular hypertonia. Eight patients had an abnormal gait; 5 were unsteady, had poor balance or fell frequently, 2 had knee flexion, patient 1 uses a wheelchair and patient 9 had wide-based walking. Sleep difficulties were reported in 7 patients, movement disorder in 2, a high pain threshold in 2, drooling in 2, stereotypies in 2 and hypothyroidism in 2. Three patients reported gastrointestinal difficulties including celiac disease in patient 3, cyclical vomiting in patient 7 and gastroesophageal reflus (resolved) – in patient 11.

### Neuroimaging for patients with MAST3 pathogenic variants

Brain MRI was reported as normal in all patients with reports available (n=5) except for patient 4 who had a smaller (left > right) anterior hippocampus. MR images for 5 patients were reviewed independently by a single pediatric neuroradiologist (S.M.); all 5 were abnormal demonstrating a small pituitary (2), thin corpus callosum (1) or both (1) (Fig 2) Patients showed no evidence of clinical features associated with a hypo-active pituitary.

**Figure 2.**
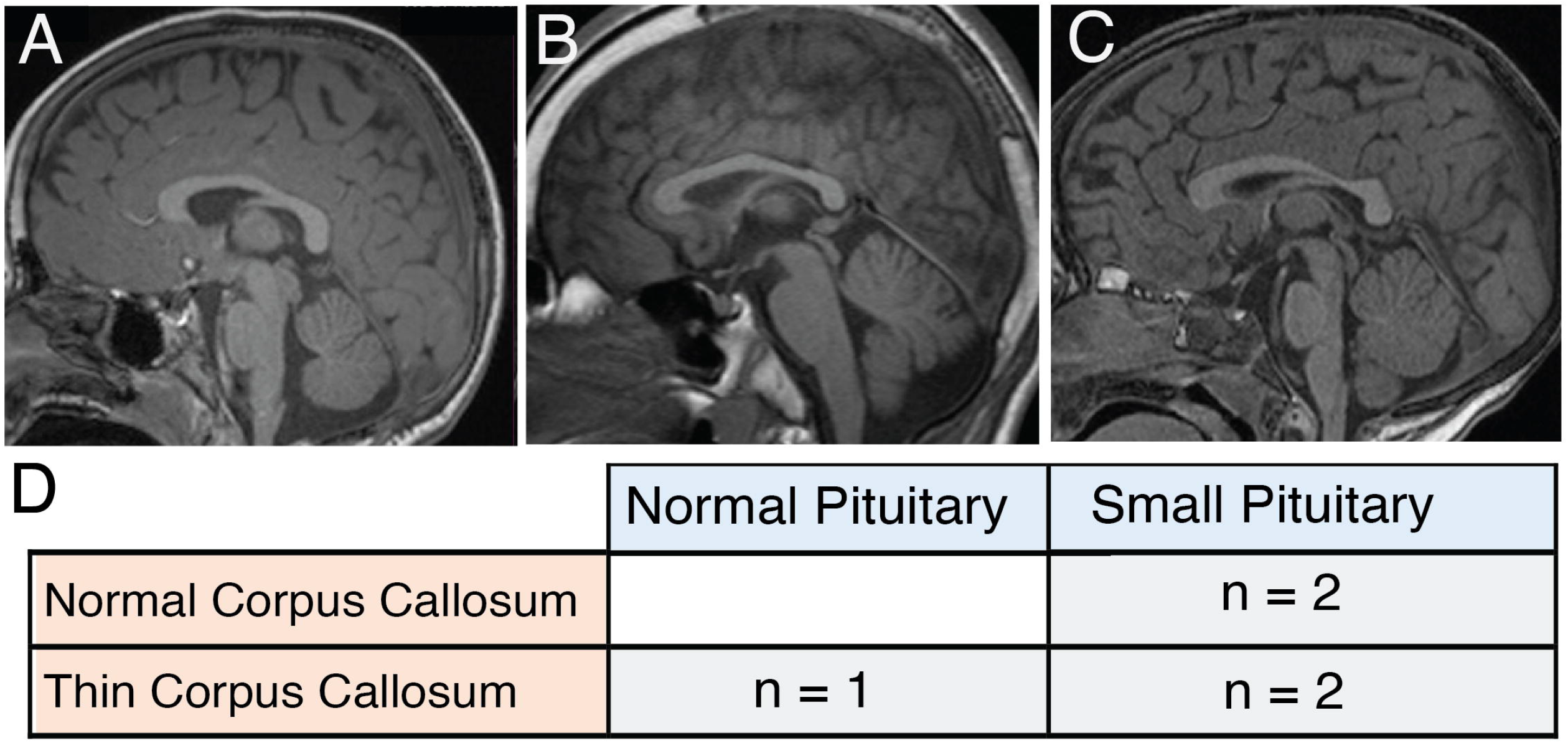
MRI changes in 5 individuals with *MAST3* variants. A-C - Sagittal FLAIR images A. Patient 7 (p.G515S) 4 years old, normal corpus callosum and small pituitary. B. Patient 11, 15 years old, thing corpus callosum and normal pituitary. C. Patient 3, 2 years 9 months old, thin corpus callosum and small pituitary. D. Table summarizing changes found in corpus callosum and pituitary gland in the 5 individuals in whom we had access to MRIs. All patients had either a thin corpus callosum and/or a small pituitary gland, but not the extensive abnormalities observed in in individuals with *MAST1* pathogenic variants.

### Expression of MAST3

We generated a heat map of *MAST1/MAST3* expression, as well as other epilepsy associated genes (*SCN1A/SCN3A*), neuronal marker genes using bulk RNA-seq data from the Brainspan atlas of the developing human brain (see list of URLs) (Fig 3A). *MAST3* expression is low over the period of human fetal neurodevelopment, with expression increasing at 26 weeks post-conception and steadily increasing postnatally (Fig 3A). Single-nuclei RNA-seq of post-mortem motor cortex from the Allen Brain Map demonstrates that *MAST3* expression is restricted to excitatory neurons in the human cortex (Fig 3B). At the protein level, MAST3 is present in a fraction of early-born (CTIP2+) neurons, and in the majority of late-born (CUX1+) excitatory neurons in 11-week-old human cerebral organoids (Fig 3C). In mouse cortex, MAST3 is present in postmitotic excitatory neurons, mostly colocalizing with CTIP2 and TBR1 at E14.5, and mainly with the upper-layer neuron marker SATB2 at E16.5 (Fig 3D).

**Figure 3.**
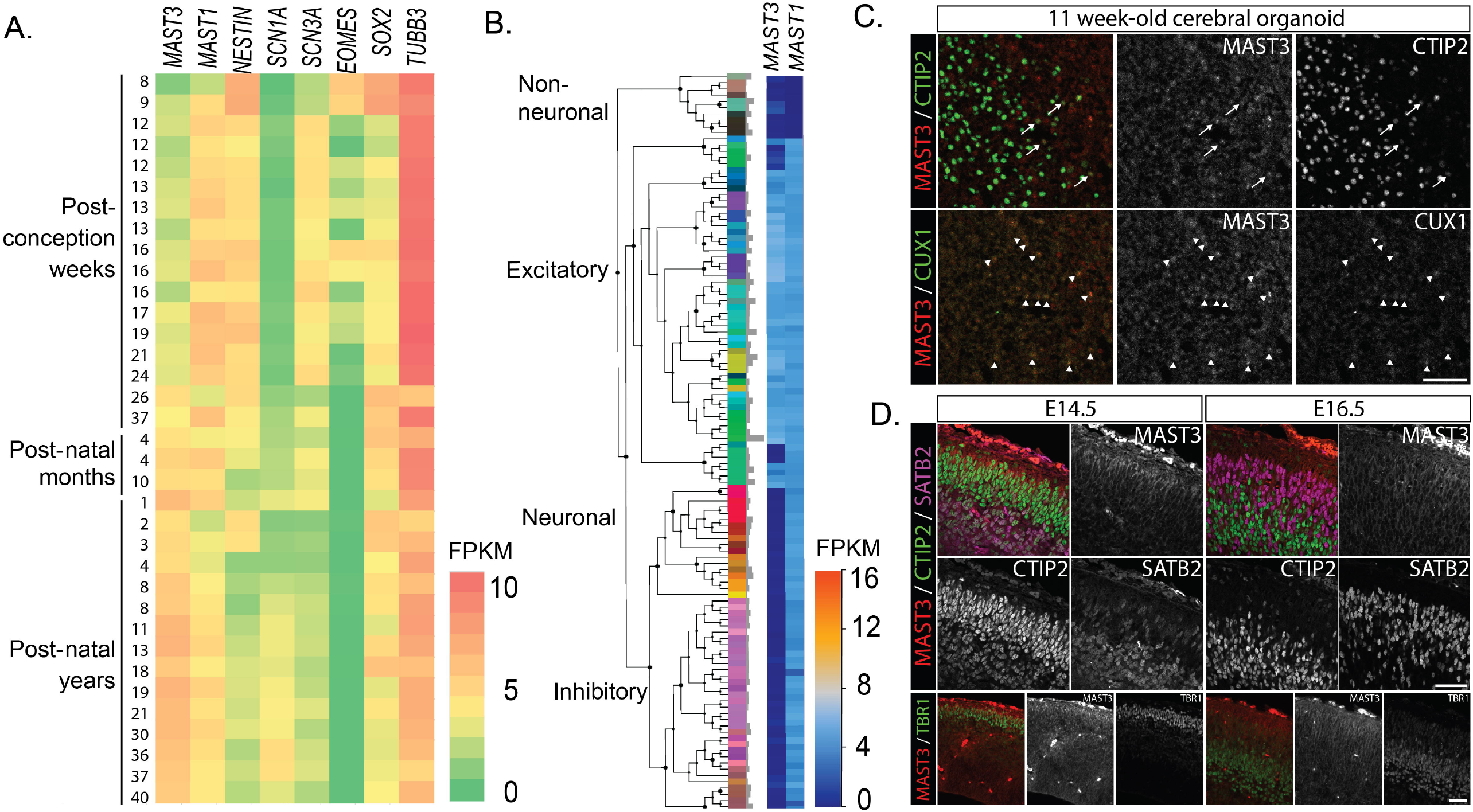
Expression of *MAST3* and *MAST1* in whole brain throughout prenatal development and postnatally. ***(A)*** Heatmap of bulk RNA-seq data from the Brainspan atlas of the developing human brain (see list of URLs). *MAST3* expression begins at about 26 weeks in prenatal development. In contrast, expression of *MAST1* is highest early in pretnatal development (9 weeks) and then is reduced throughout the lifespan. This pattern is similar in other DEE-associated genes, *SCN1A* and *SCN3A*, where only the latter is associated with malformation of cortical development ^23^. Key neuronal and developmental markers *NESTIN, EOMES, SOX2* and *TUBB3*) are also shown. FPKM = Fragments Per Kilobase of transcript per Million mapped reads. **(B)** Heatmap of single-nucleus RNA-seq data from post-mortem adult human motor cortex for *MAST3* and *MAST1* (see list of URLs). *MAST3* is expressed exclusively in excitatory neurons, while *MAST1* is expressed in both inhibitory and excitatory neurons in the cortex. **(C)** Immunofluorescence in 11-week-old cerebral organoids shows the presence of MAST3 in a fraction of excitatory CTIP2-positive neurons (upper panels, arrows) and in the majority of CUX1-positive neurons (lower panels, arrowheads). **(D)** Immunofluorescence performed in E14.5 (left panels) and E16.5 (right panels) mouse coronal brain sections confirm MAST3 expression in postmitotic upper layer neurons, co-markers evolving as cerebral development progresses. TBR1 and CTIP2 are neuronal markers specific to deep cortical layers. SATB2 is a neuronal marker for upper cortical layers. Scale bars = 50 µm.

### Assessment of kinase activity of MAST3 variants

Wild-type and four of the *MAST3* variants described in this study were expressed in HEK293T cells together with its substrate, ARPP-16 (Fig 4). A kinase-dead control (p.K396H) was also co-expressed with ARPP-16. Compared to wild-type, the expression levels of the MAST3 variants was consistently lower, with the exception of the p.V551L variant which exhibited intermediate expression (Fig 4A). As expected, wild-type MAST3 was active as demonstrated by robust phosphorylation at Ser46 of ARPP-16, while there was minimal phosphorylation by kinase-dead MAST3. Despite lower expression, the levels of Ser46 phosphorylation for each of the variants (p.G510S, p.G515S, p.L516P, p.V551L) was similar to that of wild-type MAST3. When normalized to expression levels of MAST3 protein, the activities of each variant was therefore higher than wild-type, reaching significance for the p.G510S-containing mutant (\**p* = 0.0159, one-way ANOVA with Dunnett’s posthoc test). The reason for the lower expression of the MAST3 variants is not clear but could involve decreased protein synthesis or increased protein turnover. Nevertheless, these initial studies suggest the possibility of some sort of gain-of-function phenotype for these MAST3 variants through more effective phosphorylation of ARPP-16 and possibly other substrates.

**Figure 4.**
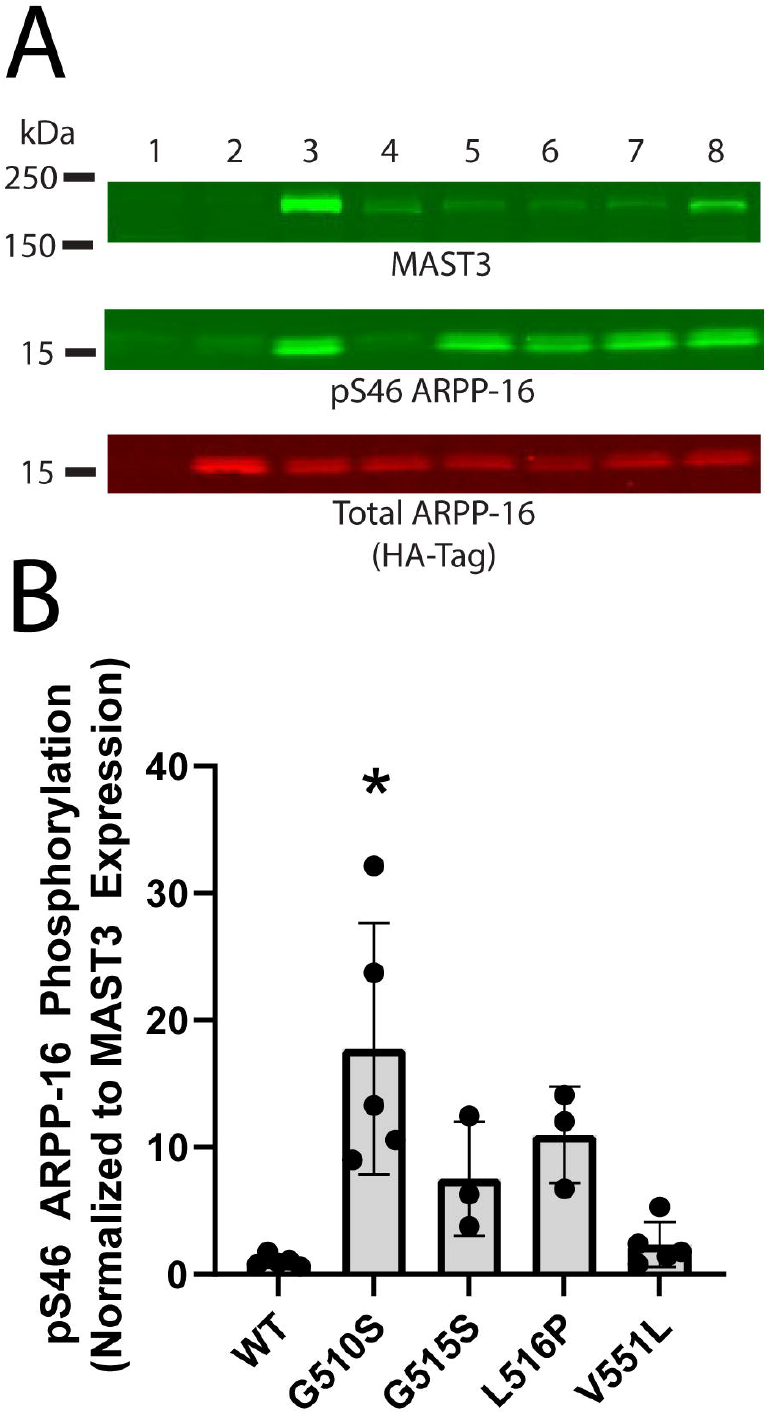
Phosphorylation of ARPP-16 at Ser46 by MAST3 kinase. ***(A)*** HA-tagged ARPP-16 alone, or together with MAST3, were expressed in HEK293T cells. Cell lysates were separated by SDS-PAGE and analyzed by immunoblotting for MAST3 protein, phospho-Ser46-ARPP-16, and HA-ARPP-16. Representative blots show that WT and all MAST3 mutant phosphorylate ARPP-16. There was minimal, background, phosphorylation of ARPP-16, as shown by the expression of ARPP-16 alone (lane 2) or when co-expressed with the dead-kinase mutant K396H (lane 4). Lanes: 1) GFP Only, 2) ARPP-16 Only, 3) Wild-Type MAST3 + ARPP-16, 4) K396H dead kinase control + ARPP-16, 5) G510S MAST3 + ARPP-16, 6) G515S MAST3 + ARPP-16, 7) L516P MAST3 + ARPP-16, 8) V551L MAST3 + ARPP-16. ***(B)*** Quantification of immunoblots revealed an increase in ARPP-16 phosphorylation when co-expressed with the G510S mutant compared to WT MAST3 (*p = 0.0159, one-way ANOVA with Dunnett’s posthoc test). Note that the GFP control, ARPP-16 alone control, and the dead kinase control quantifications were omitted from the graph due to the presence of zero values.

## Discussion

In this study, we describe de novo missense variants located in the STK domain of MAST3 as a novel cause for DEE in 11 patients. There were two recurrent variants (p.G510S and p.G515S present in 8 individuals collectively), and an additional three variants (p.R406P, p.L516P, p.V551L) located nearby ^17^. A twelfth patient harboured a genetic change outside of the STK domain; he did not have epilepsy but rather a diagnosis of ASD and additional patients and functional studies are required to interpret the pathogenicity of variants outside the STK domain. Moreover, the first variant is located within the third position of the “Asp-Phe-Gly+1” (DFG+1) domain, which determines the specificity of the target residue that will be phosphorylated and is highly conserved among protein kinases ^17, 18^.

Modulation of MAST3 by Protein Kinase A (PKA) involves phosphorylation of position 512, again close to the variants’ location in this series ^15^. Alteration of a subset of these specific variants in the STK domain resulted in lower expression in an in vitro cell culture model, but more effective phosphorylation of the MAST3 target, ARPP-16, consistent with a potential gain-of-function phenotype. Notably, a gain-of-function Gly to Ser mutation (p.G2019S) is found in an analogous position in the LRRK2 kinase and is the most common genetic cause of Parkinson’s disease^19-21^.

Patients with pathogenic variants in the STK domain of *MAST3* typically presented with DEE, including 3 patients with Dravet syndrome-like phenotype. All patients had seizure onset before 2 years of age, with 7/11 having onset before 12 months of age. Most patients had normal development prior to the onset of seizures but then experienced developmental regression or plateau which coincided with seizure onset or episodes of status epilepticus. All patients developed multiple seizure types: all 11 patients had generalized tonic clonic seizures with tonic (n=5) and focal impaired awareness seizures (n=6) also common. Triggers for seizures such as fever, illness or lack of sleep were seen in 10/11 patients. Six patients were drug-resistant whilst 3 had achieved moderate seizure control on 2-3 antiepileptic drugs and 2 had been seizure free since age 2.5-2.75 years. All patients had intellectual disability, with most (5/11) having severe impairment. Most patients developed significant comorbidities include 6 patients with autism spectrum disorder or autistic features, 8 with gait disturbances, 7 experienced sleep issues, 6 had hypotonia, 3 had dysmorphic features and 3 had gastrointestinal problems.

MAST3 is the third of the five-member MAST family of serine/threonine kinases, which share a STK and a postsynaptic density protein-95/discs large/zona occludens-1 domain (PDZ) ^5^. These genes have an overlapping but unique expression in the brain, with *MAST3* expressed predominantly in the striatum, hippocampus, and cortex ^5^. It has been studied most extensively in the rodent striatum, were the majority of *MAST3* expression is localized to medium spine projection neurons which are inhibitory GABAergic projection neurons^5^. Conversely, our transcriptomic analysis illustrates that *MAST3* expression is primarily restricted to late neurodevelopment and may be restricted to cortical excitatory neurons in humans. This pattern contrasts to MAST1, which is expressed early and at high levels in the developing brain and all cell types, including excitatory, inhibitory, and non-neuronal cells. *De novo MAST1* variants were previously described as a cause of a mega-corpus callosum syndrome with cerebellar hypoplasia ^7^. Intriguingly, this includes a p.Gly517Ser variant identified in three individuals, the same amino acid position as the MAST3 p.Gly510Ser variant described in five individuals here (Fig 1D). The authors suggest *MAST1* variants result in a dominant-negative mechanism due to reduced expression of MAST2 and MAST3 expression in a mouse model. However, here we show MAST3 variants may act in a gain-of-function manner as demonstrated by more efficient phosphorylation of its substrate, ARPP-16. Therefore, another possible explanation for the reduced expression of MAST2 and MAST3 in MAST1 knockout models is compensation for a hyperactive MAST1. Finally, the MRI abnormalities in individuals with *MAST1* pathogenic variants are extensive, while those with *MAST3* pathogenic variants we analyzed revealed very mild abnormalities (Fig 2). These discrepancies likely lie in the expression patterns of the two genes, where *MAST1* is expressed throughout development with disruption resulting in extensive malformations of cortical development while *MAST3*, expressed later and exclusively in excitatory neurons, is associated with DEE. This expression pattern is similar in the sodium channels, where *SCN3A*, which is expressed early in development, is associated with malformations of cortical development; while *SCN1A* expressed much later is associated with a DEE generally devoid of MRI findings (Fig 3A)^22, 23^. Moreover, *MAST3* expression restriction to excitatory neurons in the cortex (but not striatum) is somewhat reminiscent of *SCN8A*-associated DEE, where *SCN8A* is expressed in excitatory neurons, with gain-of-function variants leading to excessive excitatory neuron activity ^24^. We hypothesize that *MAST3* gain-of-function variants may lead to the same excessive excitatory neuronal activity, though this requires further investigation in model systems.

In both *in vitro* and *in vivo* models of striatum, MAST3 has been shown to phosphorylate ARPP-16 at Ser46 ^14, 15^, and this shifts the balance of its downstream target, protein phosphatase 2A (PP2A), from active to inactive ^25^. The *in vitro* work from this study suggests that four MAST3 variants result in a potential gain-of-function phenotype and therefore increased phosphorylation of ARPP-16. This would suggest a decrease in PP2A activity and a concomitant overall increase in phosphorylation of substrates for PP2A. Changes in PP2A have been reported in individuals with intellectual disability and developmental delays, and more specifically, the catalytic subunit of PP2A, PPP2CA (Catalytic Cα Subunit), has been implicated in patients with language delays, hypotonia, epilepsy, and brain abnormalities reminiscent of our patients ^26^.

In summary, we propose *MAST3* as a novel gene for DEE, thereby expanding the genetic landscape of DEEs. Patients with pathogenic *MAST3* variants present with DEE with normal development prior to seizure onset at < 2 years. Status epilepticus and fever-sensitive seizures are common, severe intellectual disability was seen in most as were significant comorbidities. MAST3 is the second member of MAST family of kinases to be implicated in CNS dysfunction. We propose a potential gain-of-function pathogenic mechanism that is restricted to MAST3 dysfunction in cortical excitatory neurons, reminiscent of *SCN8A*-associated epilepsy.

## URLs

Gene Matcher: https://genematcher.org/

gnomAD: https://gnomad.broadinstitute.org/ for both allele frequencies and LOEUF scores TOPMed/Bravo: https://bravo.sph.umich.edu/freeze3a/hg19/

MTR: http://mtr-viewer.mdhs.unimelb.edu.au/

Brainspan atlas of the developing human brain: https://www.brainspan.org/

Allen Brain Map single-nucleus transcriptome: https://portal.brain-map.org/atlases-and-data/rnaseq/human-m1-10x

## Acknowledgements

We would like to thank the patients and families for their participation in this research study. This work was sponsored by NIH NINDS R00NS089858 (GLC), National Health and Medical Research Council of Australia, CURE, Australian Epilepsy Research Fund, March of Dimes and NIH/NINDS (IES), the Japan Agency for Medical Research and Development (AMED) under grant numbers JP20ek0109280, JP20dm0107090, JP20ek0109301, JP20ek0109348, JP20kk0205012 (NM), JSPS KAKENHI under grant numbers JP17H01539 (NM), JP20K16862 (KI); a pilot award from the Swebilius Foundation at Yale University (ACN), NIH Common Fund, through the Office of Strategic Coordination/Office of the NIH Director under Award Numbers U01HG007690 and U01HG007530 (LHR, The content is solely the responsibility of the authors and does not necessarily represent the official views of the National Institutes of Health), KMC is an employee of GeneDx, Inc.

## Authorship contributions

ES, KRC, AGG, ACN, GLC developed and executed the study, analyzed data and wrote the manuscript, ES, EM, AS, JR, AMM, JG, ROL, ERR, BS, WGW, EJM, KI, SK, TP, CB, EHB, RHvJ, NM, LHR, KM, RG, IES, HCM, SM, LL and JJM generated and/or analysed clinical, genetic and or experimental data, all authors read and edited the manuscript.

## Conflicts of interest

GLC holds a collaborative research grant with Stoke Therapeutics. Ingrid Scheffer has served on scientific advisory boards for UCB, Eisai, GlaxoSmithKline, BioMarin, Nutricia, Rogcon, Chiesi, Encoded Therapeutics and Xenon Pharmaceuticals; has received speaker honoraria from GlaxoSmithKline, UCB, BioMarin, Biocodex and Eisai; has received funding for travel from UCB, Biocodex, GlaxoSmithKline, Biomarin and Eisai; has served as an investigator for Zogenix, Zynerba, Ultragenyx, GW Pharma, UCB, Eisai, Anavex Life Sciences, Ovid Therapeutics, Epigenyx, Encoded Therapeutics and Marinus; and has consulted for Zynerba Pharmaceuticals, Atheneum Partners, Ovid Therapeutics, Care Beyond Diagnosis, Epilepsy Consortium and UCB. She may accrue future revenue on pending patent WO61/010176 (filed: 2008): Therapeutic Compound; has a patent for SCN1A testing held by Bionomics Inc and licensed to various diagnostic companies; has a patent molecular diagnostic/theranostic target for benign familial infantile epilepsy (BFIE) [PRRT2] 2011904493 & 2012900190 and PCT/AU2012/001321 (TECH ID:2012-009). The remaining authors report no competing interests.

## Notes

### Competing Interest Statement

GLC holds a collaborative research grant with Stoke Therapeutics. IES has served on scientific advisory boards for UCB, Eisai, GlaxoSmithKline, BioMarin, Nutricia, Rogcon, Chiesi, Encoded Therapeutics and Xenon Pharmaceuticals; has received speaker honoraria from GlaxoSmithKline, UCB, BioMarin, Biocodex and Eisai; has received funding for travel from UCB, Biocodex, GlaxoSmithKline, Biomarin and Eisai; has served as an investigator for Zogenix, Zynerba, Ultragenyx, GW Pharma, UCB, Eisai, Anavex Life Sciences, Ovid Therapeutics, Epigenyx, Encoded Therapeutics and Marinus; and has consulted for Zynerba Pharmaceuticals, Atheneum Partners, Ovid Therapeutics, Care Beyond Diagnosis, Epilepsy Consortium and UCB. She may accrue future revenue on pending patent WO61/010176 (filed: 2008): Therapeutic Compound; has a patent for SCN1A testing held by Bionomics Inc and licensed to various diagnostic companies; has a patent molecular diagnostic/theranostic target for benign familial infantile epilepsy (BFIE) [PRRT2] 2011904493 & 2012900190 and PCT/AU2012/001321 (TECH ID:2012-009). The remaining authors report no competing interests.

